# Norvaline reduces blood pressure and induces diuresis in rats with inherited stress-induced arterial hypertension

**DOI:** 10.1101/678839

**Authors:** Michael A. Gilinsky, Yulia K. Polityko, Arkady L. Markel, Tatyana V. Latysheva, Abraham O. Samson, Baruh Polis, Sergey E. Naumenko

## Abstract

Growing evidence suggests that increased arginase activity affects vital bioprocesses in various systems and universally mediates the pathogenesis of numerous metabolic diseases. The adverse effects of arginase are associated with a severe decline in L-arginine bioavailability, which leads to nitric oxide synthase substrate insufficiency, uncoupling, and, eventually, to superoxide anion generation and substantial reduction of nitric oxide (NO) synthesis. In cooperation, it contributes to chronic oxidative stress and endothelial dysfunction, which might lead to hypertension and atherosclerosis.

Recent preclinical investigations point to arginase as a promising therapeutic target in ameliorating metabolic and vascular dysfunctions. In the present study, adult rats with inherited stress-induced arterial hypertension (ISIAH) were used as a model of hypertension. Wistar rats served as normotensive controls. Experimental animals were intraperitoneally administered for seven days with non-proteinogenic amino acid L-norvaline (30 mg/kg/day), which is a potent arginase inhibitor, or with the vehicle. Blood pressure (BP), body weight, and diuresis were monitored. The changes in blood and urine levels of creatinine, urea, and NO metabolites were analyzed.

We observed a significant decline in BP and induced diuresis in ISIAH rats following the treatment. The same procedure did not affect the BP of control animals. Remarkably, the treatment had no influence upon glomerular filtration rate in two experimental groups, just like the daily excretion of creatinine and urea. Conversely, NO metabolites levels were amplified in normotonic but not in hypertensive rats following the treatment.

The data indicate that L-norvaline is a potential antihypertensive agent, and deserves to be clinically investigated. Moreover, we suggest that changes in blood and urine are causally related to the effect of L-norvaline upon the BP regulation.

## 1. Introduction

Hypertension is a serious, continuously growing healthcare problem. The number of people suffering from high blood pressure (BP) has doubled over the last 40 years afflicting more than 1.13 billion people worldwide [1], including 75 million individuals in the United States alone [2]. The disease represents a leading mortality cause, with more than 7.6 million deaths per annum [3]. Moreover, there is a robust causality between devastating cardiovascular diseases, including myocardial infarction and stroke, and raised BP [3].

The hypertension etiology remains ambiguous. Recently disclosed various behavioral and genetic factors do not explicitly clarify the precise mechanisms of hypertension development. However, growing evidence indicates psychosocial factors as having an essential causative role [4]. Particular, the role of emotional stress in hypertension etiology is well-established [5]. Additionally, recent empirical data point to endothelial dysfunction and reduced nitric oxide (NO) bioactivity as the leading pathophysiological abnormalities associated with hypertension [6]. Remarkably, L-arginine supplementation reduces systemic blood pressure (BP) in some forms of experimental hypertension [7, 8] due to its direct effect upon NO synthesis and characteristic antioxidant activities, which regulate blood pressure via redox-sensitive proteins [9]. It was suggested that supplemental L-arginine is more effective in salt-sensitive hypertension than in essential hypertension [10].

Of note, L-arginine is a semi-essential amino acid, which is obtained from natural dietary sources and can be produced endogenously in various organs [11]. L-arginine plays a vital role in various physiological functions and, prominently, in maintaining vascular homeostasis [10]. L-arginine is a mutual substrate for arginase and nitric oxide synthase (NOS) (Figure 1a). Arginase is a manganese-containing enzyme that converts L-arginine into L-ornithine and urea. NOS isoforms, in turn, catalyze the production of NO and citrulline. Recent data suggest that NO-mediated vasodilation is substantially inhibited in hypertension due to an increase in arginase activity in endothelial cells (EC), which limits L-arginine availability to NOS for NO production [12].

**Figure 1:**
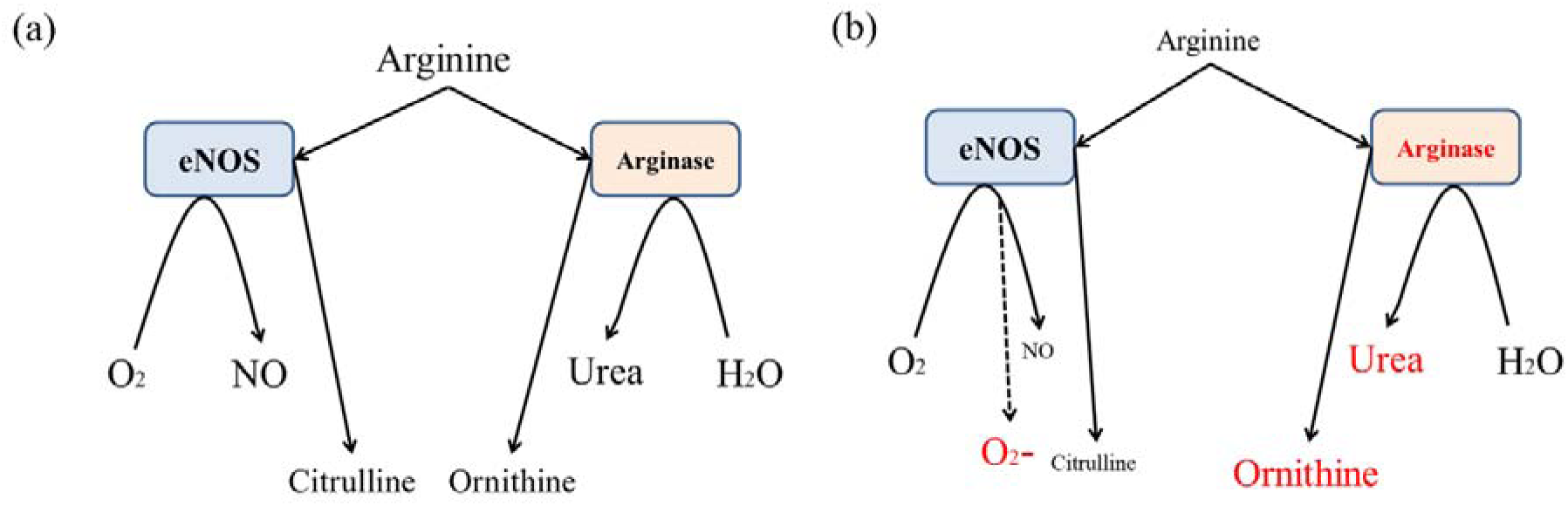
Metabolic fats of arginine in mammal cells. (a) Arginine is a mutual substrate for arginase and NOS, which are in equilibrioum in physiologic conditions. Regular coupled eNOS utilizes O_2_ and arginine to produce NO and citrulline. Arginase, in turn, converts arginine into ornithine and urea. (b) eNOS is uncoupled by substrate deficiency to produce superoxide anion rather than NO, which further diminishes NO availability.

The main effects of arginase upon BP are associated with inhibition of the NO synthesis, which is a potent vasodilator [13]. Of note, NO represents a central EC protective factor, under physiological conditions [14]; however, it becomes deleterious under oxidative stress. A decline in L-arginine bioavailability leads to endothelial NOS (eNOS) uncoupling and deflection from regular NO synthesis toward superoxide ion generation (Figure 1b) [15]. Remarkably, eNOS-deficient mice demonstrate an increased rate of atherosclerosis [16], in part, due to elevated BP [17].

Two arginase isoforms have been identified. Both types, cytoplasmic arginase I (ARG1), and mitochondrial arginase II (ARG2) have been shown to inhibit the NO production by regulating the L-arginine bioavailability [18]. While L-arginine affinity for NOS is more than 1,000 times higher than for arginase, arginase is about 1,000 times more active than NOS, which provides the equilibrium in L-arginine utilization in physiologic conditions [19]. However, the decline in substrate bioavailability and/or activation of arginase leads to a substantial shift of the balance toward ornithine synthesis (Figure 1b).

It seems that arginase activation is a conserved evolutionary reaction to various stimuli [20]. Arginase expression is inducible by catecholamines, cytokines, lipopolysaccharide, tumor necrosis factor, oxidized low-density lipoprotein (OxLDL), and hypoxia [21]. Remarkably, human and rodent aortic EC exposure to OxLDL is followed by a rapid increase in arginase activity, which is associated with ARG2 translocation from the mitochondria to the cytosol of EC [22]. Of note, hypertensive men show significantly elevated OxLDL levels compared to normotensives [23], which suggests a connection between hypertension, atherosclerosis, and arginase activation.

EC dysfunction, arising from the decline in L-arginine bioavailability and violation of the normal NOS function, (in which arginase plays a major role), leads to the development of numerous cardiovascular pathologies, and, particularly, the development of arterial hypertension. Therefore, arginase inhibition has been proposed as a therapeutic approach for the treatment of cardiovascular diseases, including hypertension [24]. Incidentally, manipulating with arginase expression levels and its activity became the goal of many studies aimed to disclose the precise mechanisms of hypertension and find ways to combat this deleterious disease.

El-Bassossy *et al*. (2013) demonstrated that arginase inhibition alleviates fructose-induced hypertension in a rat model of metabolic syndrome [25]. The authors gavaged the experimental animals with solutions of L-citrulline, L-norvaline (50 mg/kg/day), and L-ornithine for six weeks. Of note, the inhibitors utilized possess different modes of arginase inhibitory activity. The authors suggest that the effects of the arginase inhibition are directly mediated via NO signaling protection and endothelial-dependent relaxation, while indirectly associated with insulin sensitivity improvement. Another recent study by Peyton *et al*. (2018), utilizing Zucker rats as a model of obesity, evidenced a significant effect of a sustained intraperitoneal infusion of the arginase inhibitor Nω-hydroxy-nor-l-arginine (25 mg/kg/day) for four weeks on systolic BP [26]. Of note, Zucker rats display elevated BP (≈140 mmHg) from the age of 12 weeks and application of arginase inhibitor or L-arginine have a similar BP reducing effect. The same arginase inhibitor (40 mg/day) for ten weeks was used by Bagnost *et al*. (2010) in spontaneously hypertensive rats [27]. The authors speculate that the antihypertensive effect of arginase inhibition is associated with modulation of mesenteric artery reactivity, restoration of angiotensin-II-induced contraction, and acetylcholine-induced vasodilation. Another study by Pokrovskiy *et al*. (2011) convincingly demonstrated that application of an arginase inhibitor L-norvaline (10 mg/kg/day) precludes the endothelial dysfunction development in a rat model of methionine and N-nitro-L-arginine methyl ester-induced NO deficiency [28].

Of note, L-norvaline is a potent arginase inhibitor [29] and a unique compound with a broad spectrum of biological properties. It acts via negative feedback inhibition mechanism due to its structural similarity to ornithine [30] and substantially amplifies the NO production rate [31]. Moreover, in contrast to other arginase inhibitors, L-norvaline inhibits ornithine transcarbamylase (OTC), which converts ornithine to citrulline in the mitochondria [32]. The association of OTC gene polymorphisms with increased risk of hypertension has been well described [33]. Therefore, the L-norvaline application in the treatment of hypertension might be particularly beneficial.

Additionally, L-norvaline possesses anti-inflammatory properties due to its potency of inhibiting ribosomal protein S6 kinase beta-1 (S6K1) [34, 35]. S6K1is a direct downstream target of the mechanistic target of rapamycin (mTOR). Of note, growing evidence indicates that inflammation is a key factor of the hypertension pathogenesis, since endothelial dysfunction, together with oxidative stress, are evidently involved in the inflammatory cascade [36, 37]. Remarkably, inhibition of mTOR signaling pathway with rapamycin (1.5 mg/kg/day) has been shown to attenuate salt-induced hypertension in rats [38]. Accordingly, L-norvaline possesses several potentially antihypertensive modes of activity. It is worth mentioning that L-norvaline has already demonstrated a serious therapeutic potential in preclinical studies of various diseases with a clear metabolic signature. In particular, it has been suggested as a possible candidate to treat complications of diabetes mellitus [39] and Alzheimer’s disease [40].

The purpose of the present study was to investigate the therapeutic effects of L-norvaline in a rodent model of inherited stress-induced arterial hypertension (ISIAH). The ISIAH rats represent a unique model of stress-sensitive arterial hypertension [41]. The strain has been created in the Institute of Cytology and Genetics of the Siberian Branch of the Russian Academy of Sciences using Wistar rats’ background and is characterized by the genetically determined enhanced response of the neuroendocrine and renal regulatory systems to stress [42]. Consequently, the ISIAH strain is an optimal rodent model for investigation of the genetic and physiological mechanisms and pathogenesis of stress-sensitive hypertension. Particularly, we aimed to correlate the treatment-related changes in systolic BP with the blood and urine concentrations of NO metabolites in the stress-sensitive arterial hypertension.

## 2. Materials and methods

### 2.1. Animals

Studies were approved by the Biomedical Ethics Committee of the Scientific Research Institute of Physiology and Basic Medicine (SRIPBM). Permit 𝒩 <u>°</u> 7 of September 10, 2015, and conducted following the European Community Council Directive 86/609/EEC. The ISIAH rat strain was developed and bred at the animal facility of the Institute of Cytology and Genetics (Novosibirsk, Russia). Wistar rats were bred in our animal facility and served as normotensive controls. Males rats at the age of 12 weeks were used in all experiments.

### 2.2. Experimental design and treatments

Three-month-old male rats weighing about 400 g were randomly divided into four groups. Two control groups consisted of intact hypertensive ISIAH rats (n = 9) and intact normotensive Wistar rats (n = 6). Two experimental groups, with animals receiving L-norvaline treatment, consisted of ISIAH hypertensive (n=8) and Wistar normotensive (n = 6) rats.

L-norvaline (Sigma, St. Louis, MO, USA) was dissolved in isotonic saline solution. The animals were administered intraperitoneally with one ml of L-norvaline solution daily (30 mg/kg/day), or with the vehicle, at 2 PM every day for 7 days. The animals were housed individually in the metabolic cages (Italplast, Italy), which provide separation of urine and feces through the unique design of the funnel and of the separation cone. Animals had free access to balanced food and water.

Urine was collected daily at 10 AM. Blood was collected during slaughter on the eighth experimental day after decapitation on the guillotine (Figure 2). Urea in blood plasma and urine was measured using the Urea-Novo kit (Vector-Best, Novosibirsk, Russia) in accordance with the company instructions, following the method we modified. The modification consisted of using a multimodal reader TriStar LB 941 (Berthold Technologies, Germany) to detect optical density instead of using a cuvette spectrophotometer. The remaining parameters of blood and urine: creatinine, uric acid, urea nitrogen (UN) were determined by the colorimetric method using an integrated, biochemical system Dimension RxL Max (Siemens, Germany) according to the instructions of the company.

**Figure 2:**
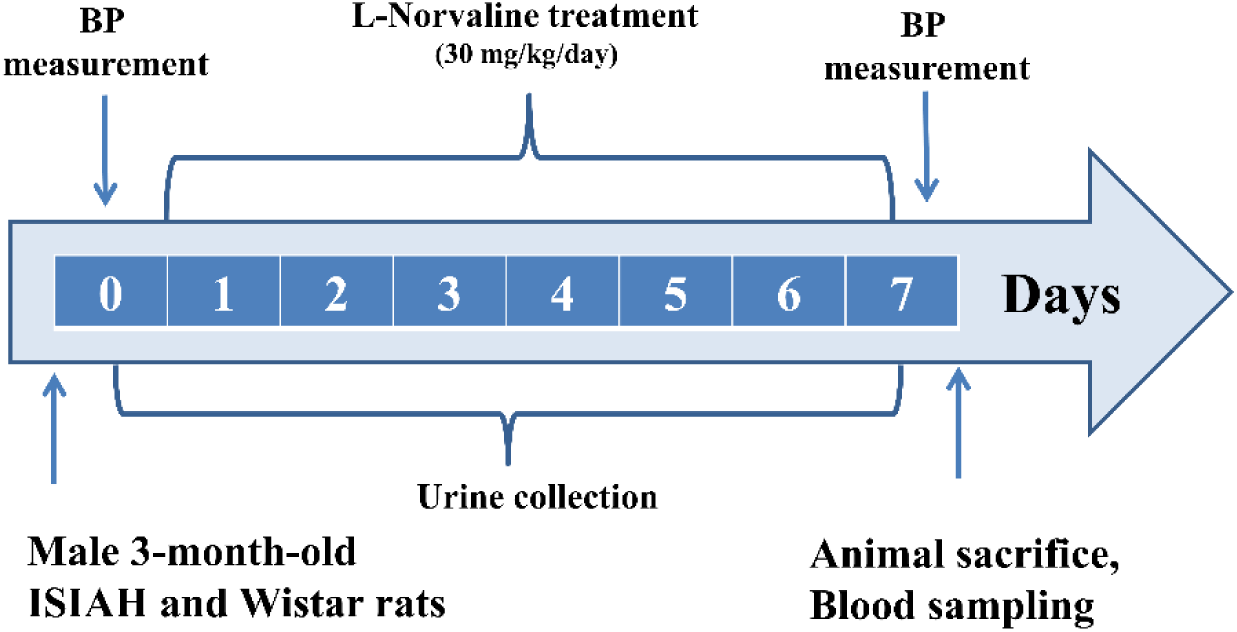
The experimental design.

### 2.3. BP measurement

For BP measurements, rats were habituated to the RR-300 restraint system (IBI Scientific, Dubuque, Iowa, USA) for 15 min/day for seven days prior to the experiment.

Blood Pressure (BP) was measured indirectly at 11-12 AM on the first and eighth days of the experiment by the same investigator. The measurements have been performed in conscious and restrained rats by the indirect tail–cuff method using a semi-automatic non-invasive BP monitoring system BIOPAC-MP system (Goleta, CA, USA) After five min of habituation in the restraint chamber, a typical series included six consequent repetitions of the automated inflation-deflation cycles. The arithmetic average value of the readings has been calculated and taken as the BP.

### 2.4. Glomerular filtration rate

Glomerular filtration rate (GFR) is a standard test utilized to evaluate the kidneys function[43]. It assesses the volume of fluid filtered from the glomerular apparatus into the Bowman’s capsule. In general practice, GFR is estimated by use of serum and urine creatinine concentrations, which is an endogenous filtration marker [44] and typically expressed in units of volume per time. In order to calculate GFR, the ratio between the urine and plasma creatinine concentrations was multiplied by the value of urine output in one minute per 100 gr of the animals’ body weight.

### 2.5. Nitrite/nitrate measurement

NO has an extremely short half-life that is about one ms in biological fluids, which substantially limits its investigations [45]. NO is scavenged by oxyhemoglobin in blood, forming nitrate, and oxidized by a copper enzyme ceruloplasmin to produce nitrite [46]. Nitrate and nitrite are relatively stable metabolites of NO, which stay stable for several hours in plasma [47]. In order to study the effect of L-norvaline treatment upon the levels of NO metabolites (nitrites and nitrates), blood and urine of the experimental animals have been subjected to the analysis. A commercially available reagent kit K1342 (Abnova, Taipei, Taiwan) has been used for the measurements. This colorimetric kit provides an accurate and convenient way to measure the total nitrate/nitrite concentration in a simple two-step process. The first step is the conversion of nitrate to nitrite using nitrate reductase. At the second stage, Griess reagent is added, which converts nitrite to a dark violet azo compound. Photometric measurement of optical density, due to this azo chromophore, accurately determines the concentration of NO_2_.

### 2.6. Statistical Analysis

Statistical analysis was conducted using SPSS version 22 (IBM, Armonk, NY, USA) for Windows. The significance was set at 95% of confidence. All the results are presented as mean with standard error. The Shapiro–Wilk test showed that the data fit a normal distribution, and Levene’s test was performed to confirm equal variance between the groups being compared. The means were compared between two groups using Student’s t-test (if appropriate) or one-way analysis of variance (ANOVA), with Tukey’s multiple comparison test used for post hoc analyses. Two-way ANOVA has been applied to check if there was an interaction between two independent variables on the dependent variable. Throughout the text and in bar plots, the variability is indicated by the standard error of the mean (SEM).

## 3. Results

### 3.1. L-norvaline effectively reduced BP in ISIAH rats

The basal levels of systolic blood pressure measured by the indirect tail-cuff method in conscious restrained rats were 123.4±1.23 (n=12) mmHg in the Wistar and 176.5±1.61 (n=15) mmHg in the ISIAH rats, which accords with our previously published data [48]. L-norvaline administration led to a substantial drop (by about 19%) in systolic BP in ISIAH rats. The main effect of the treatment was very significant F_3,28_=54.63 (p<0.0001) on the 7^th^ experimental day. The average registered BP in the ISIAH treated with L-norvaline group was 143.1±2.33 mmHg. The same treatment protocol had a minor effect upon the BP in Wistar rats. The reduction was about 13% or 16 mmHg; however, the effect was not statistically significant in this experimental group, which was evident by Tukey’s multiple comparisons post-hoc test. Two-way ANOVA analysis revealed a significant interaction between the strain and the treatment variables on BP values F1, 25=4.429 p=0.046 (Figure 3 c).

**Figure 3.**
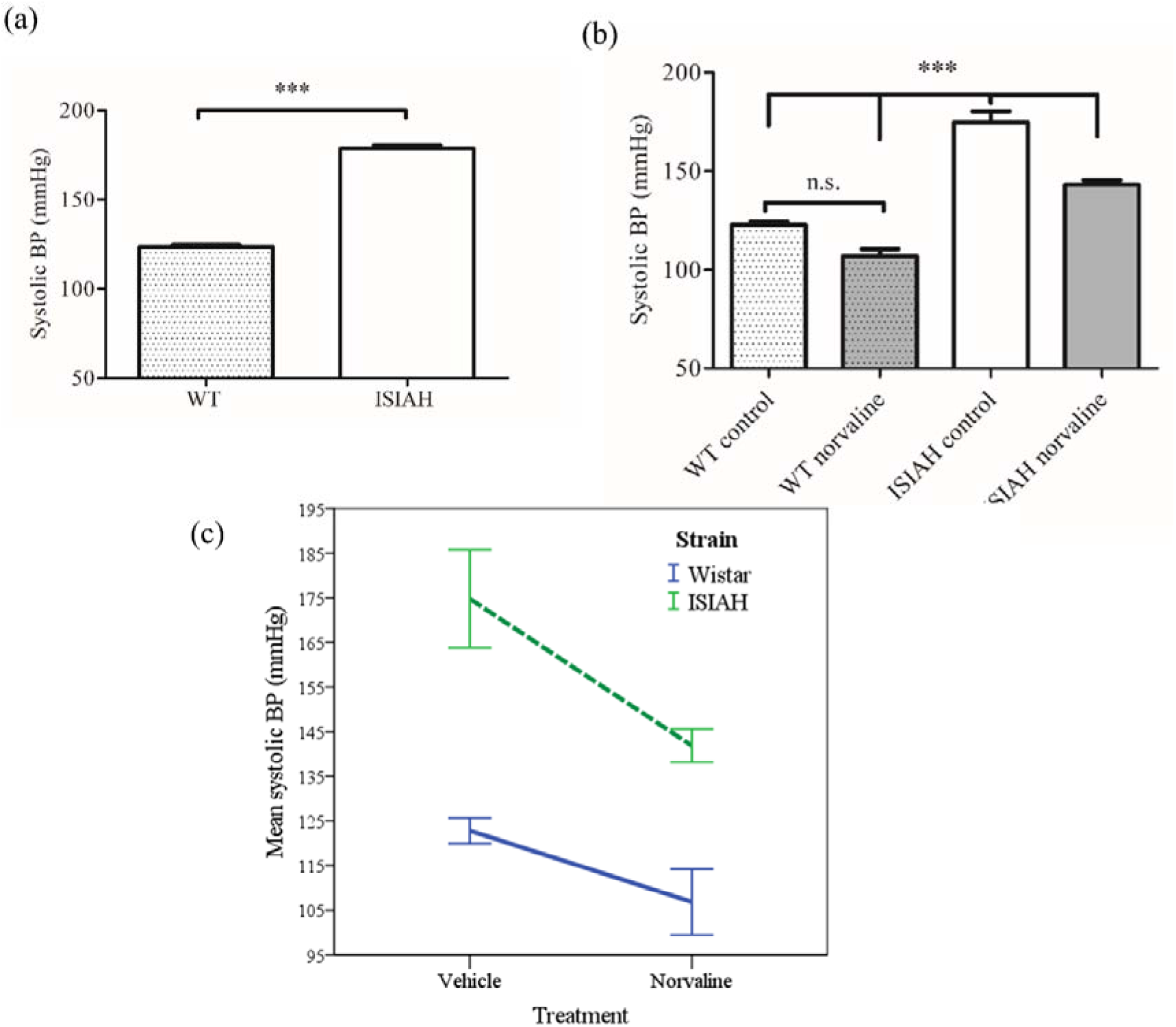
Effects of L-norvaline treatment on the systolic blood pressure (BP). (a) The mean basal levels of systolic BP measured in two experimental rat strains before treatment on day 0 (WT-Wistar rat). (b) The mean systolic BP on day 7. (c) The two-way ANOVA output representing significant differences in the graphs’ slopes. ***p < 0.001, compared with the corresponding control group values; by one-way ANOVA with Tukey’s post-hoc test. n.s –not significant. Data presented as mean ± SEM.

### 3.2. L-norvaline causes inconsequential weight loss in experimental animals

The control intact animals from two experimental strains gained about 11-12 grams on average. L-norvaline administrations for seven days led to a minor (about 5%) loss of weight, which was about 24 g in ISIAH rats and 22 g in Wistar rats on average. However, the main effect of the treatment was not statistically significant with p=0.38 and F_3,28_=1.076. Moreover, ANOVA with repeated measures did not reveal any significant effect of time upon the animals’ body weight.

### 3.3. L-norvaline treatment led to a significant decline in urine creatinine and urea concentrations in ISIAH rats but was followed by opposite metabolic effects in Wistar rats. There was no treatment-associated effect on the glomerular filtration rate in two groups

The treatment with L-norvaline led to a significant drop in the urine concentration of creatinine and urea in the hypertensive ISIAH rats compared to the intact ISIAH rats. By contrast, in the Wistar rats, the level of these urine metabolites had a clear tendency to increase (Table 1). In this regard, the daily amounts of the excreted creatinine and urea, despite interstrain differences of their concentrations, on the seventh day did not differ between the ISIAH and Wistar rats, both in control and L-norvaline treated groups. Of note, the glomerular filtration rate (GFR) in the groups of intact rats and those treated with L-norvaline was the same.

**Table 1.** The effects of L-norvaline on some blood and urine components in hypertensive (ISIAH) and normotensive (Wistar) rats on the seventh treatment day. Values are expressed as the mean ± SEM, *p<0.05.

| Rat strain | ISIAH |  | Wistar |  |
| --- | --- | --- | --- | --- |
| Type of treatment | Control | Norvaline | Control | Norvaline |
| Creatinine in urine mmol/l | 7,35±0,51 | 5,39±0,56* | 5,87±0,98 | 10,08±1,13* |
| Creatinine excretion µmol/day | 95±6,0 | 102±9,5 | 92±2,6 | 92±5,1 |
| Creatinine in plasma µmol/l | 42,4±2,5 | 38,8±1,4 | 37,2±2,3 | 37,1±1,7 |
| Glomerular filtration rate ml/min per 100 g body mass | 0,42±0,05 | 0,42±0,047 | 0,48±0,036 | 0,48±0,02 |
| Uric acid in urine µmol/l | 496±67 | 671±49 | 450±91 | 449±41 |
| Uric acid in plasma µmol/l | 34,6±7,4 | 33,8±1,3 | 36,6±2,7 | 41,6±8,3 |
| UN in urine mmol/l | 971±4,3 | 706±98,3 | 919±219,6 | 1391±111 |
| UN in plasma mmol/l | 6,79±0,29 | 6,8±0,49 | 5,97±0,57 | 5,29±0,48 |
| Urea in urine mmol/l | 1031±48,1 | 825±81,1* | 840±179,7 | 1269,2±68,5* |
| Urea excretion mmol/day | 13,63±1,24 | 17,0±2,05 | 12,5±1,15 | 12,1±1,15 |
| Urea in plasma mmol/l | 7,04±0,33 | 7,18±1,13 | 6,24±0,59 | 6,52±0,92 |

### 3.4. L-norvaline induced diuresis in the hypertonic ISIAH rats

In order to assess the effects of L-norvaline on fluid homeostasis in rats and correlate the treatment-related changes in BP with alterations in daily urine excretion and daily water consumption, we analyzed the mean water intake and urine output in the experimental groups. We observed significant strain and treatment-related differences in the daily urine output and water intake (Figure 4a,b). Of note, we did not detect any substantial differences between the groups in the basal levels of urine excretion and water intake. However, the treatment with L-norvaline was followed by a significant decline (by 17%) in the mean daily water consumption in the normotonic rats and increase (by 29%) in the mean urine excretion in the ISIAH rats. Nevertheless, the index, which reflects the general fluid balance and calculated by expressing urine output as a percentage of water intake, did not detect any significant differences between the groups, but a clear tendency to increase in hypertonic animals treated with L-norvaline (Figure 4c).

**Figure 4.**
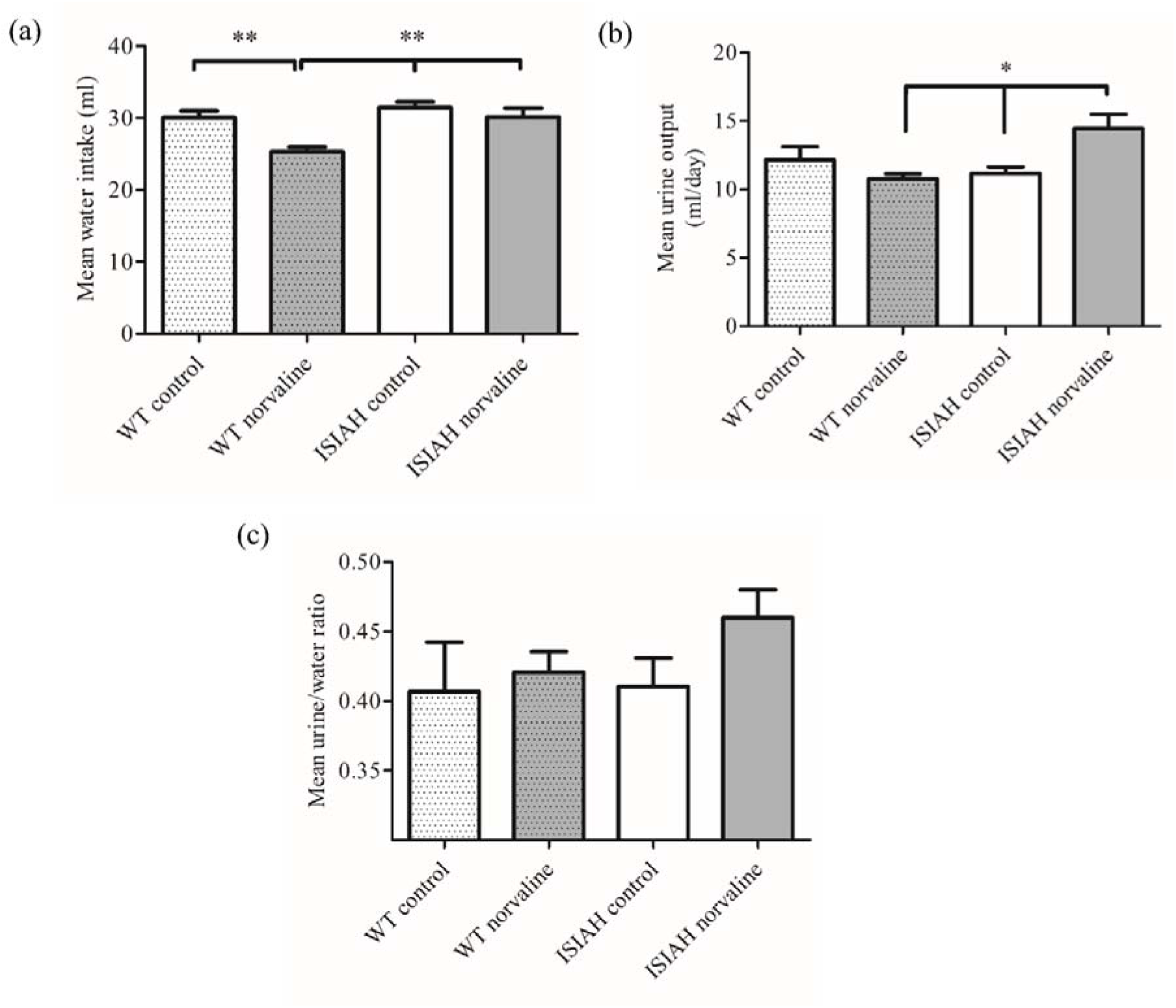
Effects of L-norvaline on fluid homeostasis. Mean water intake (a), Mean urine output (b), mean urine/water ratio (c). Water restriction was initiated immediately after dosing. Data presented as mean ± SEM. One-way ANOVA followed by Tukey’s multiple comparison test was utilized to determine significance. Significant difference indicated as **p < 0.01, *p<0.05.

### 3.5. L-norvaline amplifies the levels of NO derivatives in the plasma and urine of the wild type but not the hypertensive rats

To estimate the rates of NOS activity in the experimental animals, we quantified the urine and plasma concentrations of nitrites and nitrates, which are the stable metabolites of L-arginine-derived NO on the very last experimental day. Analysis of plasma NO metabolites (Figure 5a) revealed a significant (p<0.0001) main effect of the treatment on the blood nitrate-nitrite concentrations. Remarkably, nitrate-nitrite contents were about 34% higher in control normotensive animals than in hypertensive (1.553±0.027 vs 1.16±0.11). Moreover, the nitrite-nitrate levels increased significantly (p<0.01) by about 35% in the normotensive group following the L-norvaline treatment. On the other hand, this parameter demonstrated a moderate (about 2.5%) and statistically insignificant increase in ISIAH group. Two-way ANOVA was applied to check if there is an interaction between the strain and the treatment variables on the nitrite-nitrate concentration. The test revealed a significant main effect of the strain type and the treatment upon the dependent variable. Moreover, an interaction between strain and treatment was very significant F_1, 17_=9.282 p=0.007 (Figure 5 c).

**Figure 5.**
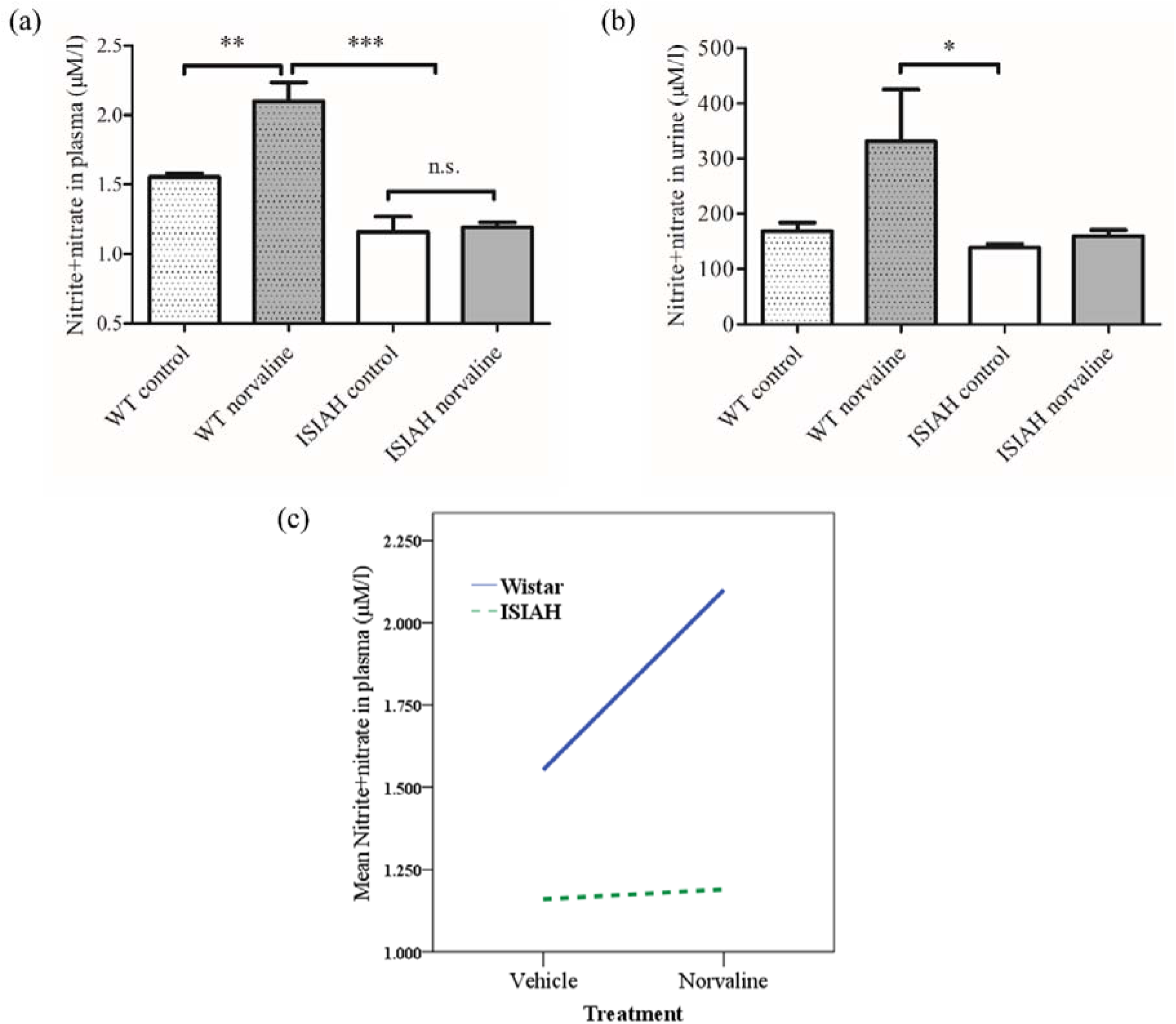
Effects of L-norvaline treatment on the levels of NO derivatives in the plasma and urine (7^th^ experimental day). (a) The mean concentrations of plasma NO metabolites. (b) The mean concentrations of urine NO metabolites. (c) The two-way ANOVA output representing significant differences in the graphs’ slopes (interaction). ***p < 0.001, **p < 0.01, *p<0.05; one-way ANOVA with Tukey’s post-hoc test. n.s. –not significant. Data presented as mean ± SEM.

Of note, the basal content of NO metabolites in the urine is more than two orders of magnitude higher than in the blood (Figure 5b). Of note, this parameter displays a substantially higher variability compared to the plasma index. Nevertheless, the plot of the urine NO derivatives levels on the last experimental day demonstrate a similar to the blood pattern, though the main effect of the treatment was significant (p=0.037). Though, an interaction between the strain type and the treatment protocol was not statistical significant p=0.142. One-way ANOVA followed by Tukey’s multiple comparison test revealed a significant difference just between Wistar treated and ISIAH control.

## 4. Discussion

There is a consensus in the literature on the crucial role of arginase in the pathological processes leading to various metabolic and cardiovascular diseases, which have a mutual pathogenesis. Up-regulation of ARG1, particularly, is prominent in myocardial infarction [49], diabetes [50], hypertension [51], and Alzheimer’s disease [35]. ARG2 activation is a leading factor of atherosclerotic vascular disease development [52], and diabetic renal injury [53]. Accordingly, arginase inhibition has been suggested to be a promising therapeutic strategy[20,54,55]. Numerous preclinical investigations testing this approach have demonstrated promising results [26]. Consequently, several clinical studies have proved that inhibiting arginase with Nω-hydroxy-nor-L-arginine significantly improves microcirculation and endothelial function in diabetic patients [56, 57].

In the present study, we demonstrated that arginase inhibition with L-norvaline reduced BP in a rodent model of inherited stress-induced arterial hypertension but had no effect upon the BP in the wild-type animals, which accords with the previously published data showing the efficacy of another arginase inhibitor N-hydroxy-nor-L-arginine in a murine model of hypertension [58]. Furthermore, we evidenced significantly decreased basal levels of NO derivatives in the blood of the hypertensive rats compared to the normotensive rats, which points to a decline in NOS activity in ISIAH rats and indicates seriously misbalanced relationships between arginase and NOS pathways, presumably due to L-arginine deprivation in this strain.

Surprisingly, we observed a shift in the L-arginine metabolism from arginase towards the NOS pathway with increased generation of NO following the L-norvaline treatment only in the wild-type group, which contrasts with the results of Bagnost *et al*. [58]. We suggest that in our rodent model of inherited stress-induced arterial hypertension, arginase is up-regulated in a much more extreme manner than in spontaneously hypertensive rats and NOS activity is more severely suppressed. Moreover, we utilized a substantially shorter treatment protocol.

Growing evidence indicates the principal role of inflammation in the pathogenesis of hypertension, which is now considered as a low-grade inflammatory condition [59]. Hypertension is empirically characterized by the presence of numerous proinflammatory cytokines in plasma and various organs. Tumor necrosis factor-α (TNF-α), above all, has been related to hypertension development and associated with the rate of renal injury. It was established that hypertensive patients exhibit higher levels of TNF-α compared to normotensive individuals [60]. Likewise, the levels of TNF-α are significantly elevated in various rodent models of hypertension. This cytokine contributes to hypertension development in Dahl salt-sensitive rats [61] and mediates hypertension in mice with angiotensin II-dependent hypertension [62]. Remarkably, the NO-deficient mouse model of hypertension is characterized by induced TNF-α generation and significant natriuretic response [63]. Shahid *et al*. (2010) elegantly demonstrated that the mice treated with a potent NOS inhibitor nitro-l-arginine exhibit substantially elevated plasma levels of TNF-α followed by an increase in mean arterial pressure, a decline in GFR and a marked escalation of sodium excretion. However, pretreatment with a TNF-α blocker blunts the effect on the sodium excretion rate [63].

In this context, the antihypertensive effect of L-norvaline might be partially attributed to its competence in inhibiting S6K1 [34]. There are strong data showing potent anti-inflammatory properties of this substance. L-norvaline treatment leads to a drop in the microglia density in the hippocampi of Alzheimer’s disease model mice, which is followed by a shift from activated to resting microglial phenotype [35]. Additionally, L-norvaline significantly reduces the brain levels of TNFα in the same murine model [40]. Therefore, its application in the models of hypertension might have a particular benefit.

In order to analyze the treatment-related changes in kidney function, we measured a list of blood and urine parameters. Even though, the examination of plasma UN and creatinine levels has no adequate sensitivity for the diagnosis of renal dysfunction, the informativity of these markers could be improved by considering the effect of body weight [64]. The GFR is one of the best indices to estimate the kidney function in health and disease [43]. We calculated this parameter per unit of the body weight and did not detect any influence of the L-norvaline treatment upon the GFR in two experimental strains, which suggests a minor role of the renal filtration in the L-norvaline-associated reduction of BP.

However, L-norvaline treatment induced diuresis in the hypertensive rats, while the same protocol had no impact upon the urine output in the Wistar rats. Previously we have demonstrated that ISIAH rats show augmented levels of tubular epithelial Na channel-alpha compared to the wild-type, which can partially explain the observed phenotype [65]. Nevertheless, we speculate that this phenomenon is more related to the treatment-associated changes in the renal NO synthesis, which are dependent upon the basal levels of the renal NOS. We have disclosed earlier a characteristic significant reduction in the juxtaglomerular NOS protein levels in the ISIAH rats [65]. This deficiency leads to renal hemodynamics impairments in this model and, presumably, plays a role in the hypertensive status formation. Accordingly, L-norvaline blocks arginase and improves substrate availability for the renal NOS, and supports NO synthesis, which normalizes the kidneys’ function, induces diuresis, and, subsequently, contributes to decreasing the elevated BP. Of note, inhaled NO has been shown to have a significant effect upon the renal function. Wraight *et al*. (2001) provided healthy middle-aged males with NO/air mixture for two hours and reported consequent increase (by about 85%) in urine output [66].

We have mentioned above that the observed increase in the ISIAH rats’ diuresis was not associated with the changes in GFR. Therefore, we speculate that the diuretic effect is related to the decrease in the sodium and water tubular reabsorption. This explanation seems plausible because NO has been shown to play a dualistic role in the regulation of renin and vasopressin secretion. Generally, NOS inhibition suppresses renin release, which indicates a stimulatory role of the L-arginine/NO pathway in the control of renin secretion [67]. However, under some conditions, and, in particular, in the models of chronic NOS inhibition, renin secretion is severely escalated [68, 69]. This phenomenon is dependent upon factors that are not clearly understood yet, and research, identifying the precise NO regulatory role in the control of renin function and urine excretion rate, still remains to be done.

## Conclusions

Our results indicate that the levels of NOS and arginase activity can be an informative biomarker to monitor the development and progression of clinical hypertension. Moreover, arginase inhibition, with L-norvaline in particular, possesses a serious therapeutic benefit due to a potential reduction of oxidative stress and inflammation, averting vascular dysfunction, and maintaining balanced physiological levels of NO.

Our findings prove significant BP-reducing and diuresis-inducing effects of L-norvaline in a murine model of stress-sensitive arterial hypertension, which suggests a potency of this agent to manage hypertension. Remarkably, a one-week-long treatment protocol has no substantial influence upon the BP in the normotensive wild type animals but leads to a meaningful BP decline in the hypertensive rats. We speculate that the observed reactions are mainly related to the immediate effects of arginase inhibition with L-norvaline upon the rate of L-arginine bioavailability, and NO synthesis in the kidney tissue, and the vessel wall in general. Moreover, we suggest that the effects of the treatment are strongly dependent on the initial parameters of the whole biological system, which are responsible for the regulation of renal circulation rate, and BP maintenance that are expressively misbalanced in the adult ISIAH rats [42].

Additionally, we suggest that it would be of value to investigate the contribution of the different arginase isoforms to the development of stress-induced arterial hypertension. The transgenic animals with up-regulated and/or deleted arginase genes might represent attractive research models.

## Authors’ Contribution

Michael Gilinsky, Arkady Markel, Abraham O. Samson designed the experiments. Michael Gilinsky and Yulia Polityko performed the experiments. Tatyana Latysheva, Yulia Polityko, Baruh Polis, Sergey Naumenko analyzed the data. Michael Gilinsky and Baruh Polis wrote the manuscript. Arkady Markel, Sergey Naumenko, Baruh Polis and Abraham O. Samson advised, developed experimental methods and edited manuscript.

## Acknowledgments

Russian Foundation for Basic Research supported this work with grant 𝒩<u>°</u> 17-04-00916.

